# An amyloidosis-associated polymorphism does not alter LECT2 stability *in vitro*

**DOI:** 10.1101/2022.03.01.482540

**Authors:** Liudmila Belonogov, Paris E. Taylor, Sherry Wong, Gareth J. Morgan

## Abstract

Amyloid fibrils formed from leukocyte chemotactic factor 2 (LECT2), a secreted human cytokine, are associated with kidney failure in the disease amyloid LECT2 (ALECT2) amyloidosis. This rare disease was recognized in 2008 and has a variable prevalence worldwide. The mechanisms which lead to ALECT2 fibril deposition are not known and there are no treatments other than kidney transplant. The LECT2 gene harbors a single nucleotide polymorphism that leads to either a valine or isoleucine residue at position 40 of the mature protein. Most of the individuals diagnosed with ALECT2 amyloidosis are homozygous for valine at this position, which led us to hypothesize that the valine-containing variant of LECT2 protein is less stable and more prone to aggregation than the isoleucine-containing variant. Here, we investigate the structure, stability and aggregation of both variants of recombinant LECT2. Both variants have similar structures in solution; unfold in similar concentrations of urea; and aggregate at similar rates under native-like conditions, forming structures that bind to thioflavin T. Chelation of the structural zinc ion destabilizes both variants to a similar extent, and increases the rate at which they aggregate. We do not observe a consistent difference in stability or aggregation between the variants of LECT2, so we suggest that the presence of the valine residue at position 40 does not determine whether an individual is at increased risk of ALECT2 amyloidosis.

## Introduction

Deposition of the protein leukocyte chemotactic factor 2 (LECT2) as insoluble amyloid fibrils can lead to progressive kidney damage (Benson, 2010; Benson et al., 2008; Larsen et al., 2014; Murphy et al., 2010). The resulting disease is known as amyloid LECT2 (ALECT2) amyloidosis (Benson et al., 2020). First identified in 2008, ALECT2 amyloidosis is recognized as a rare but serious condition, with variable incidence worldwide (Benson et al., 2008; Larsen et al., 2016; Mereuta et al., 2014; Rezk et al., 2018). Recent surveys suggest that ALECT2 fibrils are among the most common causes of renal or hepatic amyloidosis, amongst patients whose amyloid requires characterization from an organ biopsy (Mereuta et al., 2014). No treatments are available, and patients eventually progress to renal failure and may require transplant. Little is known about the causes of the disease.

LECT2 is a 133 amino acid cytokine secreted mainly by the liver (Yamagoe et al., 1996, 1998). The LECT2 gene encodes a 20-residue, N-terminal signal peptide which is removed before secretion. LECT2’s functions are not fully understood, but it has been linked to immune and inflammatory signaling and lipid metabolism, primarily in the liver (Lan et al., 2014; Lu et al., 2016; Shen et al., 2016). This signaling may be involved in the pathogenesis of diabetes, liver fibrosis and cancer (Chen et al., 2014; Lan et al., 2014; Xu et al., 2019). ALECT2 fibrils are composed of the full-length mature protein (Benson et al., 2008). The secreted protein has a metalloproteinase-like structure, in which a zinc ion is coordinated by two histidine residues and one aspartic acid residue (Okumura et al., 2013; Zheng et al., 2016). Three disulfide bonds between cysteine residues are present in the folded state. Despite its protease-like structure, recombinant LECT2 appears not to possess endoprotease or exoprotease activity *in vitro* (Zheng et al., 2016). Instead, the apparent substrate binding site may be associated with binding to receptors including Tie1 (Xu et al., 2019) and MET (Chen et al., 2014).

Several forms of systemic amyloidosis are associated with inherited mutations that destabilize the native state of the precursor protein (Benson et al., 2020; Rowczenio et al., 2014; Sekijima et al., 2005). Although individuals with ALECT2 amyloidosis do not carry an analogous mutation, most or perhaps all have a shared genotype. Position 172 of the chromosomal gene is a known single-nucleotide polymorphism site (SNP rs31517) (Kameoka et al., 2000). On the coding strand, this nucleotide can be adenine or guanine, coding for isoleucine or valine, respectively, at position 40 of the mature LECT2 protein, with population-wide allele frequencies of 0.62 and 0.38, respectively. Individuals with ALECT2 amyloidosis studied to date are often homozygous for G172, and hence V40 (Mereuta et al., 2014; Murphy et al., 2010). The frequency of V40 homozygosity in the population is around 0.15, so this variant cannot be sufficient to cause disease. However, it may represent a risk factor for ALECT2 amyloidosis.

We hypothesized that mutation of isoleucine 40 to valine (I40V) in LECT2 would destabilize the native structure and/or increase its propensity to aggregate, thus predisposing individuals with this polymorphism to ALECT2 amyloidosis. The effect of a mutation is highly dependent on the structural context: the variants I107V and V122I in the protein transthyretin are both associated with inherited forms of amyloidosis (Rowczenio et al., 2014). The crystal structure of LECT2 (Zheng et al., 2016) shows V40 buried in the hydrophobic core of the protein (Figure 1A). The additional methyl group of an Ile residue could potentially stabilize or destabilize such a structure. Since removal of a buried hydrophobic moiety is generally associated with destabilization, we predict that V40 is the less stable form, which would be consistent with an increased propensity to form amyloid fibrils. To test this hypothesis, we expressed and purified I40 and V40 variants of recombinant LECT2, and investigated their structure, stability and propensity to aggregate *in vitro*.

**Figure 1:**
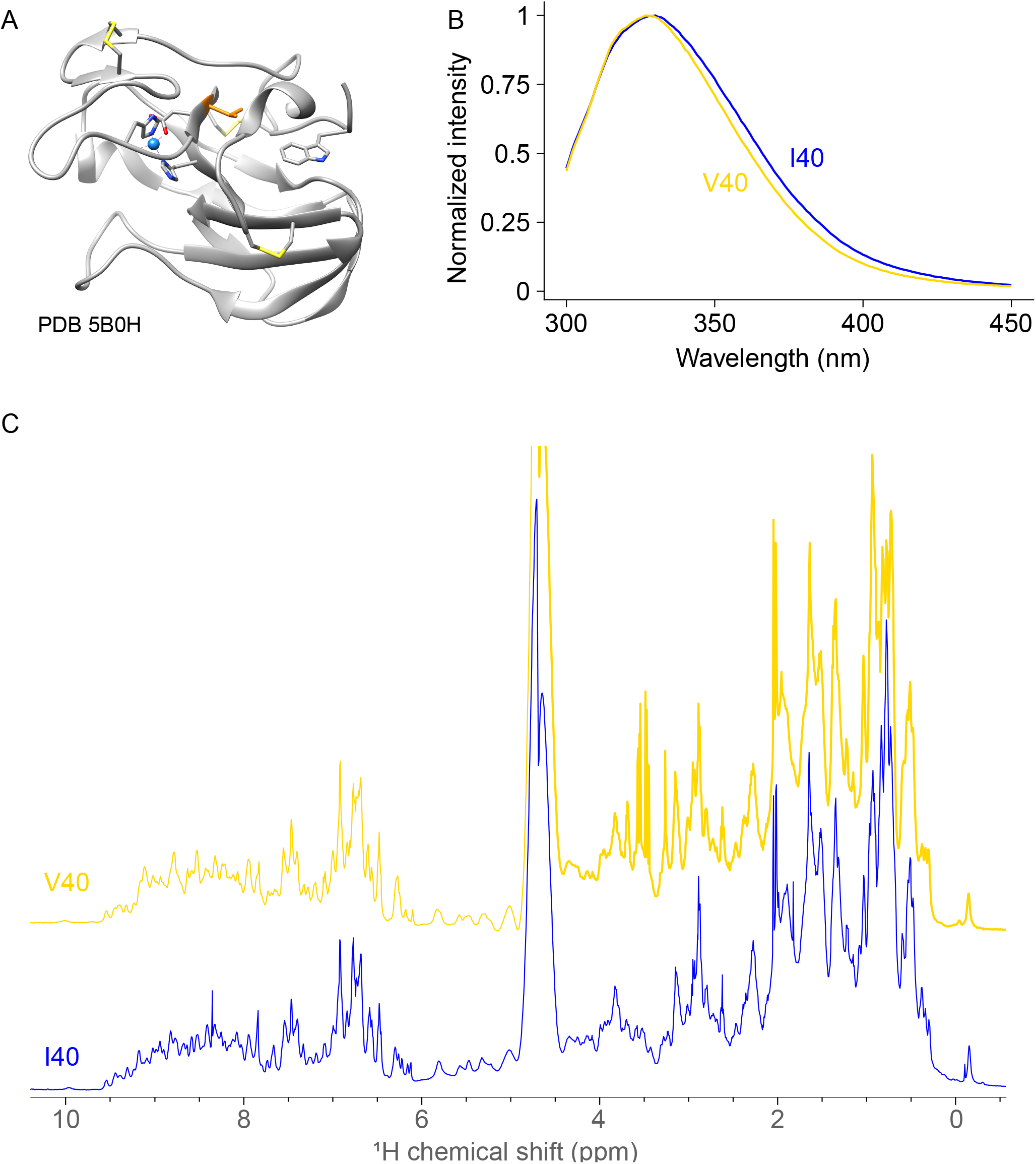
LECT2 variants form similar structures *in vitro*. A) Crystal structure of the LECT2 V40 variant, PDB code 5B0H. The Zn^2+^ ion is shown as a cyan sphere, and structurally relevant residue are shown as sticks. Valine 40 is highlighted in orange. Image prepared using UCSF Chimera. B) Fluorescence emission spectra (λ_ex_ = 280 nm) of LECT2 V40 (gold) and I40 (blue) are similar. Intensities are normalized to the peak value to account for differences in concentration. C) ^1^H NMR spectra of LECT2 V40 (gold) and I40 (blue) are similar.

## Materials and Methods

### Materials

All reagents were from Sigma (MO, USA) or Fisher Scientific (NH, USA) unless otherwise noted.

### Protein expression and purification

A synthetic DNA fragment encoding LECT2 was purchased from Twist Biosciences and cloned into a pET-11a expression plasmid (EMD, MA, USA). The V40I mutation was created using a Q5 Site Directed Mutagenesis kit (New England Biolabs, MA, USA). The resulting plasmids, pET-11-LECT2-V40 and pET-11-LECT2-I40, have been deposited with Addgene under accession codes 183031 and 183032. Despite LECT2’s internal disulfide bonds, we observed good expression from BL21 (DE3) *E. coli* without the need for refolding. We verified expression and purification by immunoblotting with an anti-LECT2 antibody (#MAB722, R&D Systems, MN, USA). Cells were grown in terrific broth media at 37 °C, cooled to 12 °C and induced with 0.1 mM IPTG, then incubated overnight to allow protein expression. Cells were harvested by centrifugation and stored at −20 °C. Cell pellets were lysed by sonication and cell debris removed by centrifugation and filtration. The LECT2 proteins were purified by cation exchange chromatography using an SP sepharose column (Cytiva Life Sciences, MA, USA) at pH 6, eluting with a 0-1 M gradient of NaCl. Size exclusion chromatography on a Superdex 75 column (Cytiva), equilibrated with phosphate buffered saline (PBS, 1.5 mM KH2PO4, 8.1 mM Na2HPO4, 2.7 mM KCl, 138 mM NaCl, pH 7.4), was used to remove remaining impurities and ensure that the purified LECT2 proteins were monomeric. Elution volumes were consistent with monomeric LECT2 (250 ml from a 320 ml column) but this was not verified further.

### Spectroscopy

Fluorescence spectra were acquired on a Horiba Fluorolog instrument at 25 °C in 50 mM sodium phosphate buffer, pH 7. Excitation was at 280 nm and emission measured between 300 and 450 nm. Spectra were corrected by subtracting the spectrum of the appropriate buffer measured under identical conditions. The wavelength at maximum intensity (λmax) was determined by fitting to a spline curve in R. To monitor changes in fluorescence over the urea titration, we first calculated the average wavelength <λ>, or center of spectral mass (Royer et al., 1993), was calculated according to the formula:

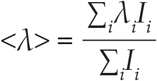

where *λi* and *Ii* are the wavelength and intensity, respectively, at each data point i. These data were then normalized to the 0M and 8M urea values to allow comparisons between variants, according to the formula:

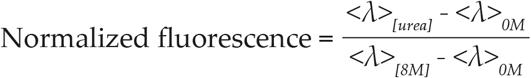

where <λ>[*x*] is the average wavelength at a molar urea concentration of *x*.

Samples for nuclear magnetic resonance (NMR) spectra were concentrated to approximately 200 μM and exchanged into in 10 mM sodium phosphate buffer containing 10% D_2_O, pH 7. One-dimensional proton spectra were recorded, using a Watergate sequence for solvent suppression (Piotto et al., 1992), at 25 °C on a Brucker Avance spectrometer equipped with a cryoprobe, operating at a proton frequency of 500 MHz. Spectra were analyzed using Brucker TopSpin software.

### Stability measurements

LECT2 proteins (5 μM) were incubated at 25 °C in 50 mM sodium phosphate buffer, pH 7 containing progressively increasing concentrations of urea. For measurements of *apo*-state stability, 1 mM ethylene diamine tetraacetic acid (EDTA) was added to the solutions to chelate the Zn^2+^ ion. For measurements of the Zn-bound state stability, an additional 10 μM ZnCl_2_ was added to minimize loss of Zn from otherwise folded protein. After incubation at 25 °C for 2 hours, tryptophan fluorescence emission spectra were recorded. Excitation was at 280 nm and emission measured between 300 and 450 nm. Data were exported to R and processed using the Tidyverse suite of tools.

### Amyloid formation assays

LECT2 variants were diluted to 10 μM in PBS containing 1 μM thioflavin T (ThT) and either 10 μM ZnCl_2_ or 1 mM EDTA. 200 μl aliquots (n=6 independent replicates per plate) were added to the wells of black 96-well plates with clear bottoms (Corning #3631). The plates were sealed with film and covered by a lid, then incubated at 37 °C before fluorescence readings were taken. ThT fluorescence was measured using a Molecular Devices Spectramax M5 platereader (λ_em_ = 440 nm, λ_ex_ = 480 nm), reading through the bottom of the plate to avoid evaporation. After an initial reading, plates were incubated at 37 °C, shaking at 1000 rpm on an orbital plate shaker. Fluorescence was measured at intervals to track aggregation. Data were exported to R and processed using the Tidyverse suite of tools.

## Results

We first asked whether the two LECT2 variants form similar tertiary structures. We expressed the I40 and V40 variants in *E. coli* and purified them by column chromatography. We expressed untagged, mature LECT2, lacking its signal sequence, to avoid potential interactions between a hexahistidine tag (Ito et al., 2003) and the integral Zn^2+^ ion. Both variants expressed sufficiently well in the cytosol of BL21 (DE3) *E. coli* to provide enough recombinant protein for the experiments that follow, although we have observed significant variability in yield between batches.

LECT2 contains a single tryptophan residue that is buried in the hydrophobic core of the protein, according to the crystal structure of LECT2 V40 (Figure 1A). Intrinsic tryptophan fluorescence spectra (λ_ex_ = 280 nm) of both LECT2 I40 and LECT2 V40 are similar, with λmax values of 329 nm and 328 nm, respectively (Figure 1B). We also analyzed the conformations of the proteins by solution nuclear magnetic resonance (NMR). Both variants have similar ^1^H NMR spectra, with dispersed peaks in the amide regions and upfield-shifted methyl peaks (Figure 1C). These features are consistent with folded structures and indicate that the secondary and tertiary structures of the LECT2 variants are similar.

Native LECT2 contains a bound Zn^2+^ ion that may stabilize its structure. To verify that an ion is present in the coordination site, we measured fluorescence and NMR spectra in the presence of excess EDTA, a divalent cation chelator. We observed a red-shift in the fluorescence spectra (λ_ex_ = 280 nm), consistent with an increased solvent exposure of the single tryptophan residue (Figure 2A). The NMR spectral linewidths increased following addition of EDTA, consistent with increased conformational heterogeneity (Figure 2B). The strongest peak below 0 ppm, which probably originates from a methyl group close in space to an aromatic moiety, is significantly reduced in intensity. However, the overall shapes of the spectra were similar. The amide regions of the NMR spectra remain dispersed, consistent with retention of hydrogen bonded secondary structural elements. These observations are consistent with *apo*-LECT2 populating an dynamic ensemble of structures that includes native-like conformations. The presence of the Zn^2+^ ion therefore appears to promote the native-like conformation.Based on these data, we conclude that both recombinant LECT2 variants fold to a structure similar to that observed in the crystal, stabilized by Zn^2+^.

**Figure 2:**
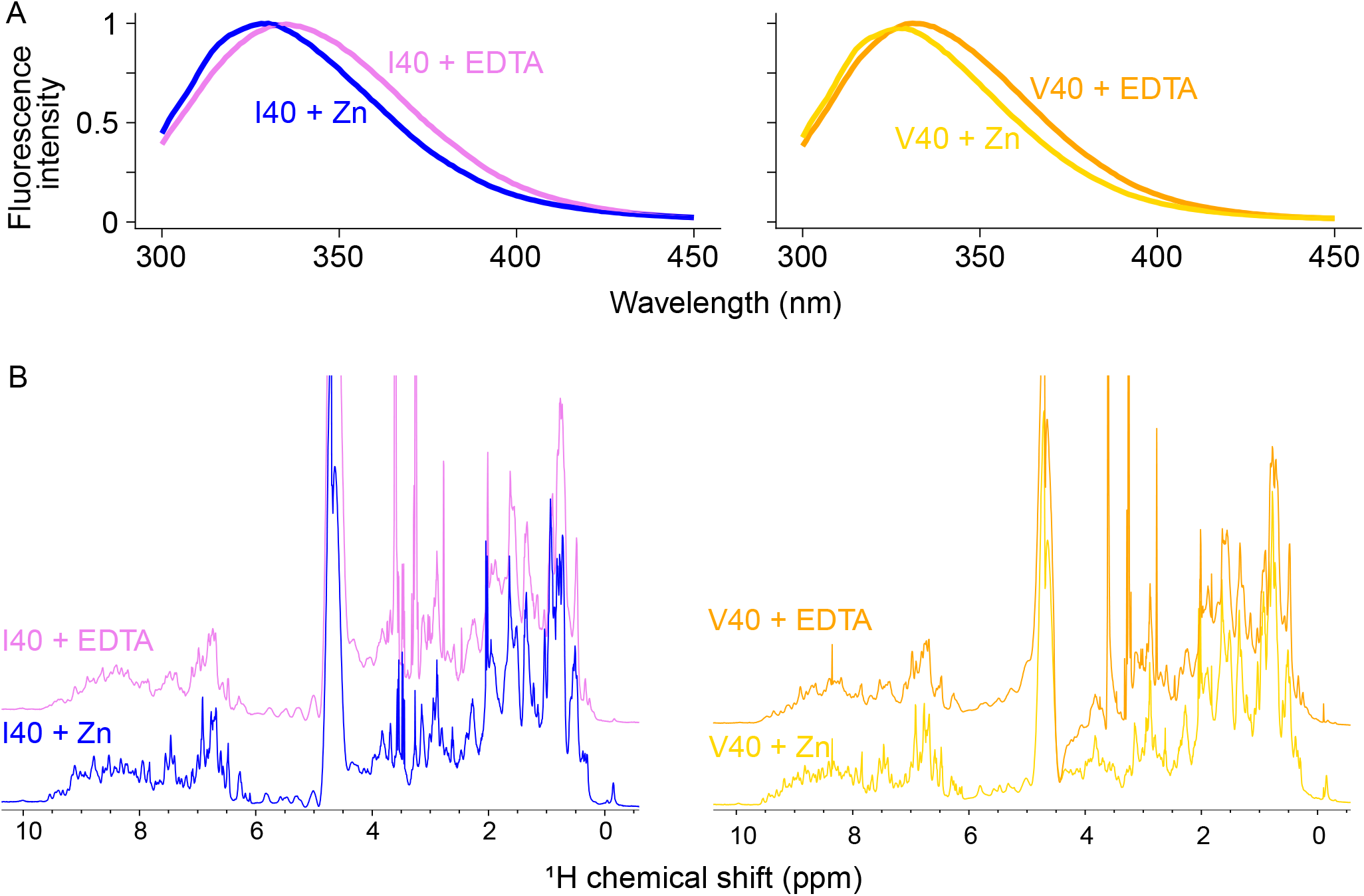
Removal of the bound Zn^2+^ ion alters the structure of LECT2. Left, 5 μM LECT2 I40 in the presence (magenta) or absence (blue) of 1 mM EDTA. Right, 5 μM LECT2 V40 in the presence (orange) or absence (gold) of 1 mM EDTA. A) Fluorescence emission spectra (λ_ex_ = 280 nm). B) H NMR spectra.

To ask whether the native structures of LECT2 I40 and LECT2 V40 are similarly stable, we measured denaturation by urea titration, following the conformation of the protein by tryprophan fluorescence. Addition of 8 M urea to LECT2 results in a change in the tryptophan emission spectrum consistent with solvent exposure of this residue (Figure 3A). Titration of urea (Figure 3B) revealed that both variants unfold at indistinguishable urea concentrations (5 +/- 0.2 M at 25 °C, pH 7). Addition of 1 mM EDTA to the unfolding titrations significantly reduced the concentration of urea required for unfolding, consistent with reduced stability of the proteins. However, the behavior of the I40 and V40 *apo* states are very similar (Figure 3C).

**Figure 3:**
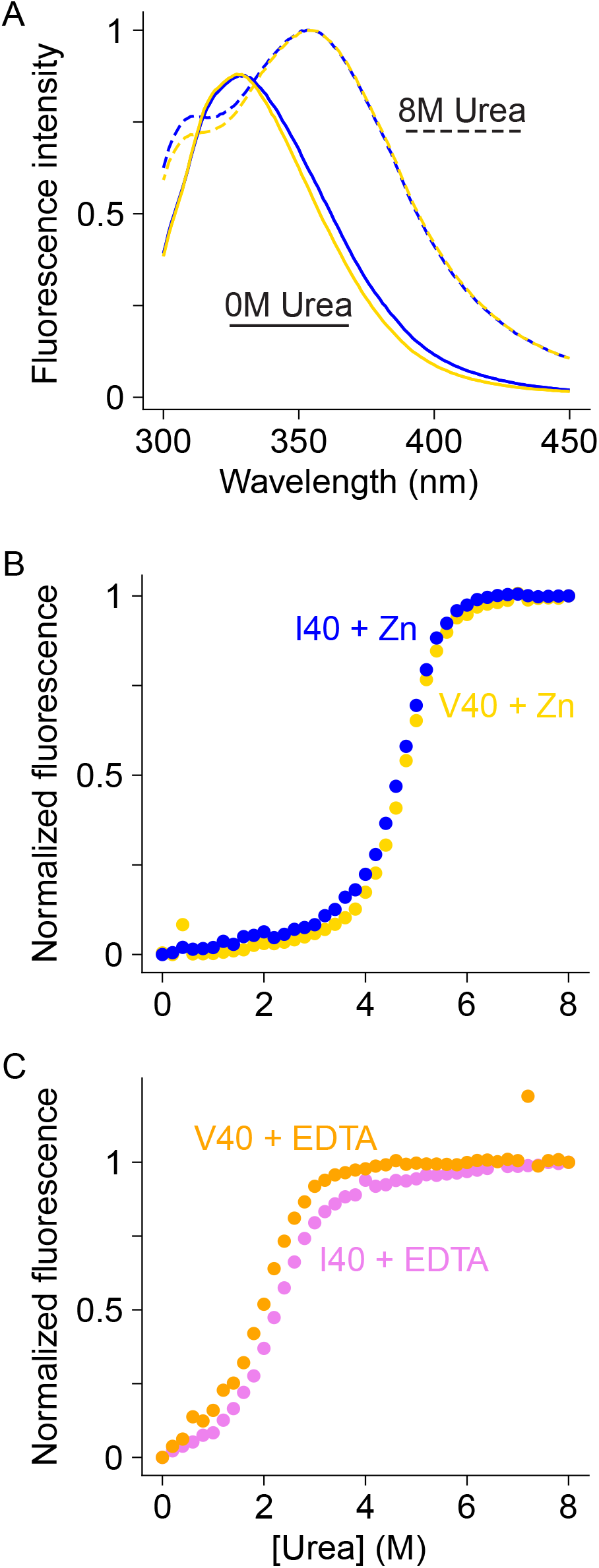
LECT2 I40 and V40 variants have similar stability. A) Fluorescence emission spectra (λex = 280 nm) of LECT2 I40 (blue) and V40 (gold) in the absence (solid lines) presence (dashed lines) of 8M urea. B) Urea titrations in the presence of 10 μM ZnCl_2_. C) Urea titrations in the presence of 1 mM EDTA.

We next asked whether LECT2 could aggregate *in vitro* to form amyloid-like fibrils. Incubation with shaking (1000 rpm) at 37 °C, pH 7.4 in microwell plates led to formation of aggregate species that bound to the amyloid-sensitive dye thioflavin T (ThT) within 24 h. The rates at which fibrils were formed was similar in both variants (Figure 4), although the kinetic profiles are different. The amplitudes and apparent rates of change of ThT fluorescence increase are lower for V40 than for I40, but the lag phase for V40 may be shorter than that for I40. There is significant variation between the rates of amyloid formation in individual wells on the same plate, which reduces our ability to identify differences between the two LECT2 variants. Chelation of Zn^2+^ by EDTA leads to more rapid and more reproducible aggregation kinetics, consistent with the observation that Zn^2+^ chelation destabilizes the protein (Figure 4). Again, there is no consistent difference between the two variants of LECT2. Further experiments may resolve differences between the two variants, but there is no clear evidence that one variant aggregates more readily than the other.

**Figure 4:**
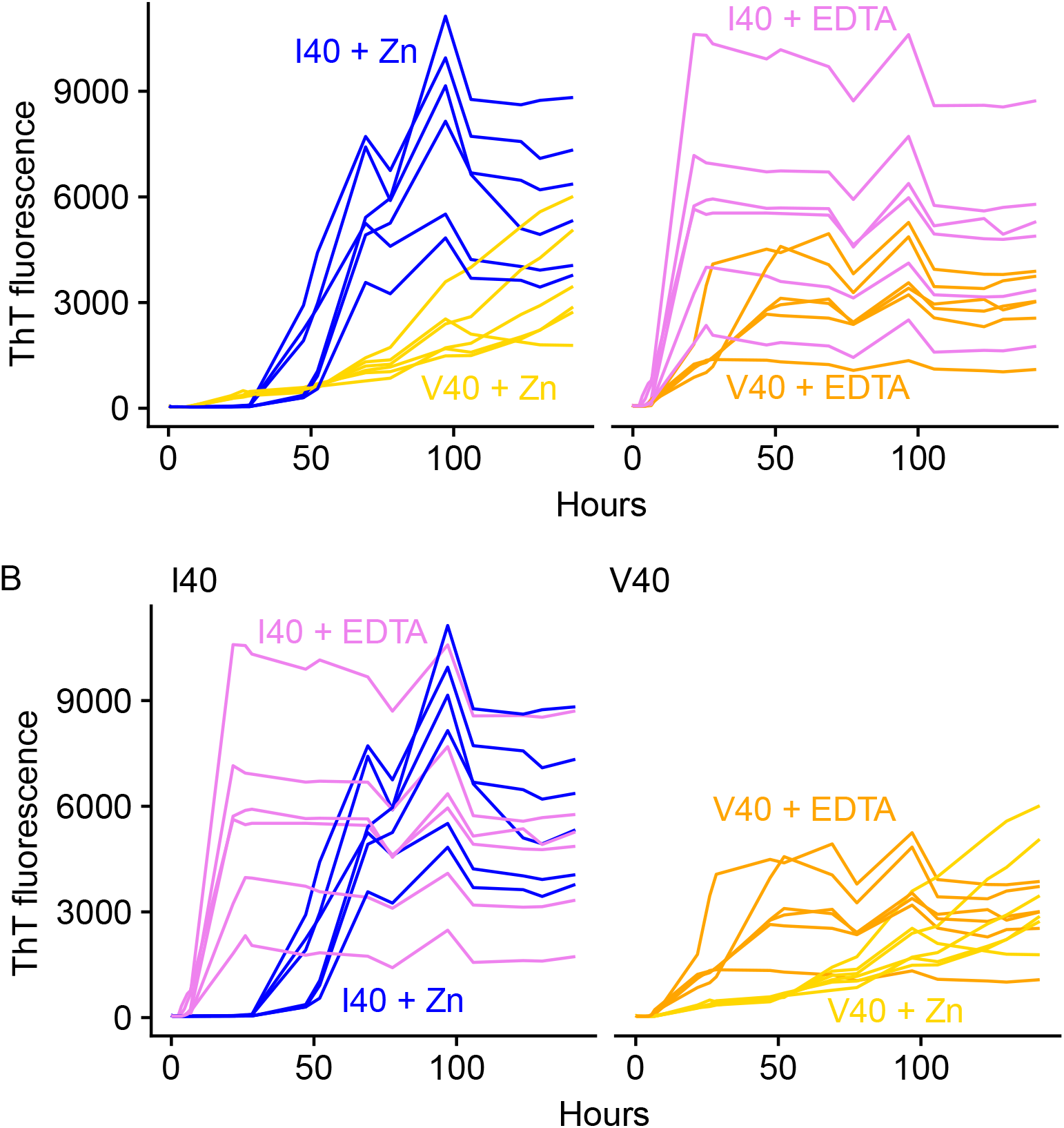
LECT2 I40 and V40 variants aggregate to form species that bind to thioflavin T (ThT). Fluorescence traces from six wells per condition are shown. A) Comparison of LECT2 I40 (blue and magenta) with V40 (gold and orange) in the presence of 10 μM ZnCl2 (left) or 1 mM EDTA (right). B) The same data plotted to show the effect of Zn^2+^ chelation by EDTA on each variant.

Based on these experiments, we conclude that the LECT2 I40V polymorphism has only a very small effect on the protein’s stability and aggregation *in vitro*. Although we cannot rule out an effect *in vivo* that is not recapitulated by our recombinant protein, we do not see strong evidence that the V40 variant is more amyloidogenic.

## Discussion

Although the V40 variant of LECT2 is associated with ALECT2 amyloidosis, we did not find evidence that this polymorphism leads to destabilization of the protein, or acceleration of aggregation *in vitro*. Both recombinant LECT2 variants form similar structures *in vitro*, and both have similar stabilities, as measured by urea titration and resistance to aggregation. In contrast, removal of the structural Zn^2+^ ion substantially destabilizes both proteins and makes them more prone to aggregation. Loss of the native fold of LECT2 appears to be required for amyloid formation, as has been observed for other proteins.

Although there are small differences between the variants, we do not consider them sufficiently robust to reject the null hypothesis that the I40V polymorphism has little effect on stability or amyloid formation *in vitro*. Further experiments are planned, but this work was significantly disrupted by the pandemic restrictions in 2020 and is currently suspended. Given the lack of published biophysical data on LECT2, we feel that these data may be useful to researchers and share them on that basis, but we caution that the data should be regarded as preliminary.

The genotype at the polymorphism at SNP rs31517 appears not to be directly causative for amyloidosis via the mechanisms described for inherited forms of ATTR amyloidosis or other familial amyloid diseases. There may be factors in patients, such as altered protein-protein interactions or post-translational modifications that we are not able to test in this model. However, the I-to-V mutation is relatively conservative, apparently not greatly altering the biochemical or biophysical properties of the LECT2 protein. It is possible that homozygous individuals have a greater risk of amyloidosis, because the two variants may not readily form copolymers, such that homozygous individuals have a greater concentration of the amyloid-forming protein. A similar scenario appears to determine risk of prion diseases (Palmer et al., 1991). The V40 variant encoded by SNP rs31517 is more common in individuals who identify as Hispanic or Latino in the US, who are over-represented among ALECT2 amyloidosis patients, so this polymorphism may also co-segregate with another risk factor (Larsen et al., 2014). We suggest that the guanine allele of SNP rs31517 is unlikely to be a predictive risk factor for ALECT2 amyloidosis, and that individuals who are not homozygous at this position may still be at risk of amyloidosis.

The stabilizing effect of the coordinated Zn^2+^ ion in LECT2 may also be involved in amyloidosis. Hereditary mutations in the gelsolin gene that destabilize one of its Ca^2+^-chelating repeat domains and reduce that domain’s affinity for Ca^2+^ have been implicated in amyloid gelsolin (AGel) amyloidosis (Solomon et al., 2012). In AGel amyloidosis, increased susceptibility to proteolysis by furin or other proteases leads to production of amyloidogenic fragments (Huff et al., 2003; Solomon et al., 2009). Although proteolysis is not known to play a role in ALECT2 amyloidosis, similar destabilization involving disruption of metal binding could be involved. Alterations in metal homeostasis could also potentially contribute to pathology.

Our observations lead us to hypothesize that destabilization of either variant of LECT2 in patients could contribute to ALECT2 amyloidosis. Although there is little difference in stability between the two variants, removal of the structural Zn^2+^ ion by EDTA chelation substantially reduces the stability of LECT2 and renders it more prone to aggregation. Analogous to the pathology of ATTR amyloidosis, this implies that pharmacologic stabilization of native LECT2 protein could reduce amyloid deposition and perhaps benefit patients (Bulawa et al., 2012). The lack of a biophysical rationale for the association of the V40 variant with disease highlights that little is known about pathogenesis of this rare but increasingly recognized disease.

## Acknowledgements

This work was supported by the Charles Brown Research Fund of the Boston University Amyloidosis Center. PET was supported by the Boston University Summer Training as Research Scholars program, which is funded by NLHBI Award R25HL118693. We thank Dr. C. James McKnight and the Department of Physiology and Biophysics at Boston University School of Medicine for use of the NMR spectrometer, which was purchased with funds from NIH Shared Instrument Grant S10OD011941.

